# Phase-locking of neural activity to the envelope of speech in the delta frequency band reflects differences between word lists and sentences

**DOI:** 10.1101/2023.05.31.543025

**Authors:** Yousef Mohammadi, Carina Graversen, Jan Østergaard, Ole Kaeseler Andersen, Tobias Reichenbach

## Abstract

The envelope of a speech signal is tracked by neural activity in the cerebral cortex. The cortical tracking occurs mainly in two frequency bands, theta (4 - 8 Hz) and delta band (1 - 4 Hz). Tracking in the faster theta band has been mostly associated with lower-level acoustic processing, such as the parsing of syllables, whereas the slower tracking in the delta band relates to higher-level linguistic information of words and word sequences. However, much regarding the more specific association between cortical tracking and acoustic as well as linguistic processing remains to be uncovered. Here we recorded electroencephalographic (EEG) responses to both meaningful sentences as well as random word lists in different levels of signal-to-noise ratios (SNRs) that lead to different levels of speech comprehension as well as listening effort. We then related the neural signals to the acoustic stimuli by computing the phase-locking value (PLV) between the EEG recordings and the speech envelope. We found that the PLV in the delta band increases with increasing SNR for sentences but not for the random word lists, showing that the PLV in this frequency band reflects linguistic information. When attempting to disentangle the effects of SNR, speech comprehension, and listening effort, we observed a trend that the PLV in the delta band might reflect listening effort rather than the other two variables, although the effect was not statistically significant. In summary, our study shows that the PLV in the delta band reflects linguistic information and might be related to listening effort.

## 1. Introduction

When listening to speech, cortical activity tracks the low-frequency amplitude modulation (envelope) of the speech signal (Ding et al., 2015; Ding & Simon, 2013; Giraud & Poeppel, 2012; Pasley et al., 2012). This cortical tracking plays a functional role in speech processing. In multi-talker listening situations, for instance, the cortical tracking of target speech is modulated by selective attention (Kerlin et al., 2010; O’Sullivan et al., 2015; Rimmele et al., 2015) and by the intelligibility of distractor speech (Dai et al., 2022): the cortical tracking increases when attention is focused on the target stream and decreases when distractor intelligibility increases. Moreover, stimulating the auditory cortex through the transcranial alternating current with signals that are derived from the speech envelope can modulate and even enhance the comprehension of speech in background noise (Keshavarzi et al., 2020; Riecke et al., 2018; Wilsch et al., 2018; Zoefel et al., 2018).

The cortical tracking of speech in different frequency bands presumably relates to different aspects of speech perception. In particular, tracking in the delta frequency band (1-4 Hz) was found to correspond to word-level, phrasal, and acoustic prosodic features, and the theta band (4-8 Hz) to syllabic features (McHaney et al., 2021; Peelle et al., 2013). However, the precise roles of speech tracking for lower-level acoustic as well as higher-level linguistic processes are still debated. Some studies argued that neural speech tracking is restricted to the processing of acoustical cues (Howard & Poeppel, 2010; Millman, Johnson, & Prendergast, 2015; Nourski et al., 2009), while others suggested that non-sensory linguistic information, such as semantic and syntactic information, increased cortical tracking of speech envelope (Ahissar et al., 2001; Luo & Poeppel, 2007; Meyer et at., 2018; Peelle & Davis, 2012; Peelle et al., 2013).

The impact of linguistic information on cortical speech tracking emerged, for instance, from studies that reported increased cortical tracking of connected speech compared to control stimuli, such as noise-vocoded unintelligible speech (Peelle et al., 2013; Rimmele et al., 2015), time-reversed speech (Gross et al., 2013; Molinaro & Lizarazu, 2018), or speech in a foreign language (Ding et al., 2015). In particular, Ding et al. (2015) demonstrated that during listening to connected speech, cortical activity tracks the linguistic structures at different hierarchical levels such as words, phrases, and sentences if the participants understood the speech. The cortical responses to the sentential and phrasal information were, however, absent in participants who did not understand the language, despite the acoustic stimuli being the same.

To which extent background noise impacts the cortical tracking of the speech envelope remains debated. Ding & Simon (2013) showed that the neural tracking of target speech is relatively insensitive to the level of background noise, but others reported that target speech tracking significantly decreased as the level of noise progressively increased (Ghinst et al., 2019; Petersen et al., 2017; Ghinst et al., 2016). The latter effect might emerge because noise reduces speech intelligibility through the energetic or informational masking of target speech (Wang & Xu, 2021) and makes speech recognition challenging by affecting the segregation and selection of acoustic speech streams from background noise.

Dimitrijevic et al. (2019) showed that cortical speech tracking quantified by speech-brain coherence (in the range of 2-5 Hz) is related to the level of listening effort that participants expend in a digits-in-noise task. In particular, low speech-brain coherence was associated with higher listening effort. They also found that the correct identification of digits was related to increased speech-brain coherence. Decruy et al. (2020) also tested the hypothesis that speech tracking is modulated by the amount of listening effort in a speech-in-noise task with different levels of signal-to-noise ratios but found no significant association of the latter to delta-band speech tracking. However, they reported that self-reported listening effort could explain a small part of inter-subject variability in the theta-band cortical tracking of the speech envelope. Specifically, they reported that when speech understanding was above 50%, envelope tracking decreased with increased effort. On the other hand, when speech comprehension was below 50%, the opposite relation was observed, i.e. higher envelope tracking was associated with enhanced effort.

In this study, we further examined the effect of background noise level and linguistic information on the neural tracking of speech. In particular, we sought to disassociate the influence of linguistic content on the putative relation between listening effort and cortical speech tracking. To this end, we measured behavioral responses as well as EEG to two types of speech stimuli that differed in their linguistic content: meaningful sentences and random word lists. Both types of stimuli were presented in different levels of background noise to yield variability in speech comprehension and listening effort. We used the phase-locking value (PLV) to quantify speech-brain phase locking and investigated the temporal dynamics of this value at different frequencies.

## 2. Materials and Methods

### 2.1. Dataset

The present study used previously collected EEG datasets (Mohammadi et al., 2023). The recording of this dataset is described in detail below.

### 2.2. Participants

Participants were 32 healthy native Danish speakers (two left-handed, 13 females, mean age 24 ± 3 years) with normal hearing and no history of neurological, psychiatric illness, or use of psychotropic medication. All gave written informed consent and were compensated financially for their participation. The number of participants was selected based on previous studies on the effect of listening effort on neural responses (Dimitrijevic et al. 2019, Decruy et al. 2020). The study was conducted in accordance with the Declaration of Helsinki and was approved by the ethics committee of Northern Jutland, Denmark (N-20200061).

### 2.3. Stimuli

Two types of speech stimuli with different linguistic information were used: sentences and random word lists. Sentences were obtained from the Dantale II database (Wagener, Josvassen, & Ardenkjær, 2003). This database consists of 150 sentences. Each sentence was generated by a random combination of the alternatives of a base list. The base list consisted of ten sentences, each containing a subject, verb, numeral, adjective, and object with the same syntactical structure but semantically unpredictable (e.g., in English: "Ulla owns five red jackets"). All sentences were recorded by a female native Danish speaker at a sampling rate of 44.1 kHz. The duration of the sentences varied from 1.85 s to 2.52 s (2.22 s ± 0.12 s).

Random word lists were created that had neither sentence-level semantic content nor syntactic structure. Each sentence of the base list was split into five words, yielding 50 different words. A natural pause after each word was kept by selecting the duration of individual words from the beginning of the given word up to the beginning of the next word. Word lists were created by randomly combining five words from the list of 50 words (e.g., in English: "rings get nine fourteen sold"). The duration of all word lists was between 1.58 s and 2.71 s (2.20 s ± 0.16 s), comparable to that of the sentences, thus avoiding a confound of different durations between the two types of stimuli that could occur otherwise (Kolozsvári et al., 2021).

Both types of speech material were read with normal sentence prosody. We verified that there were accordingly no significant differences in prosody between sentences and word lists (Figure 2). In particular, the onset time of the words was indistinguishable between the two types of speech material (see below for statistics). For example, the mean onset time of the second word in the sentences was 451 ms (after speech onset), and in the word lists, it was 443 ms. There were significant differences between sentences and word lists in the average envelope in the time intervals of 0 - 258 ms and 1340 – 1720 ms (p < 0.001, analyzed by cluster-based permutation t-test). Although not ideal, the prosodic difference between sentences with syntactic structure and word lists without structure is inherent and acceptable, given that word onsets are not significantly different. No significant difference was observed in the envelope power spectrum (p > 0.05) between the sentences and the word lists.

**Figure 1.**
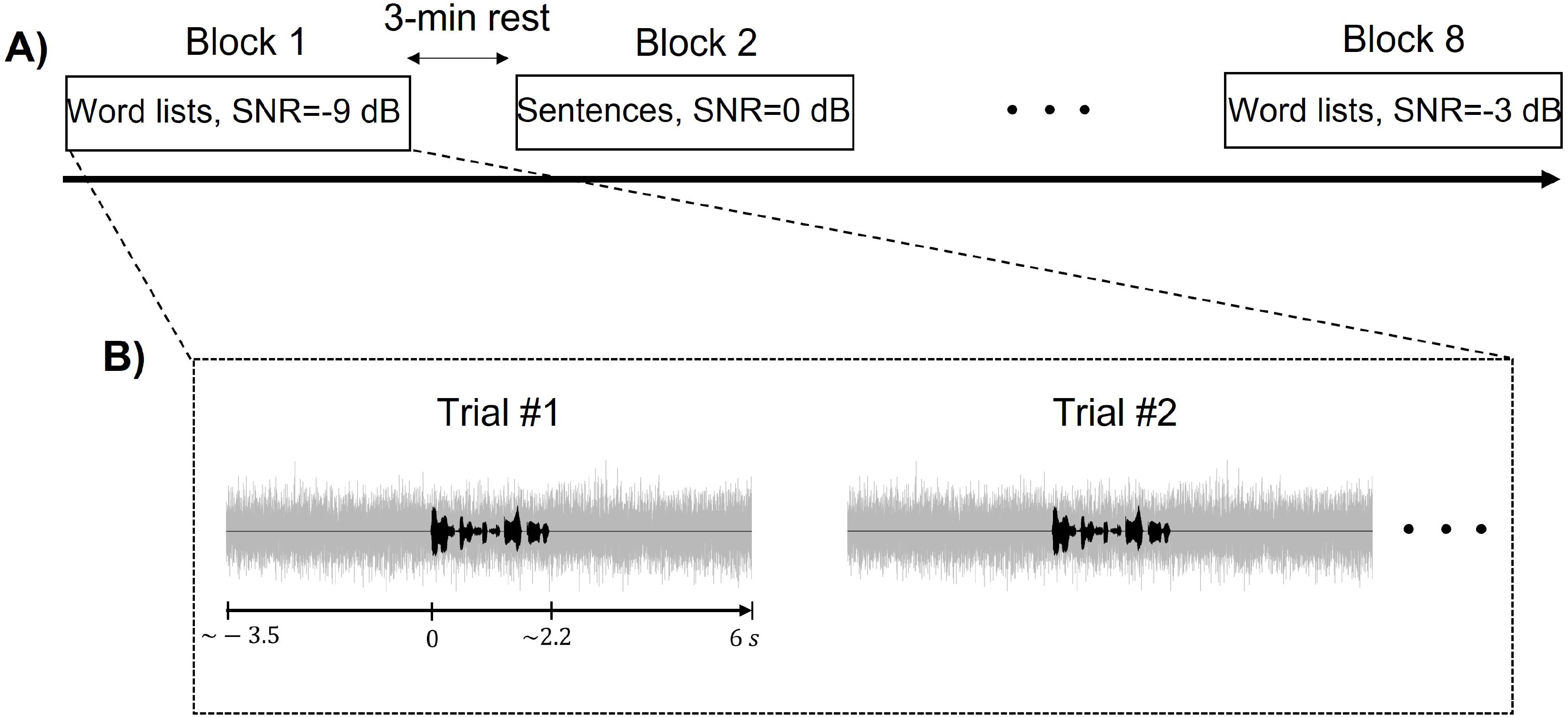
Experimental procedure. (**A**) Speech was presented in different blocks. Each block was randomly assigned one of four signal-to-noise ratios (SNRs) (−9 *dB*, −6 *dB*, −3 dB, and 0 dB) and one of two speech types (sentences and word lists), resulting in eight conditions. After each block, participants had a three-minute rest. (**B**) Each trial began with background noise in which the speech material was embedded.

**Figure 2.**
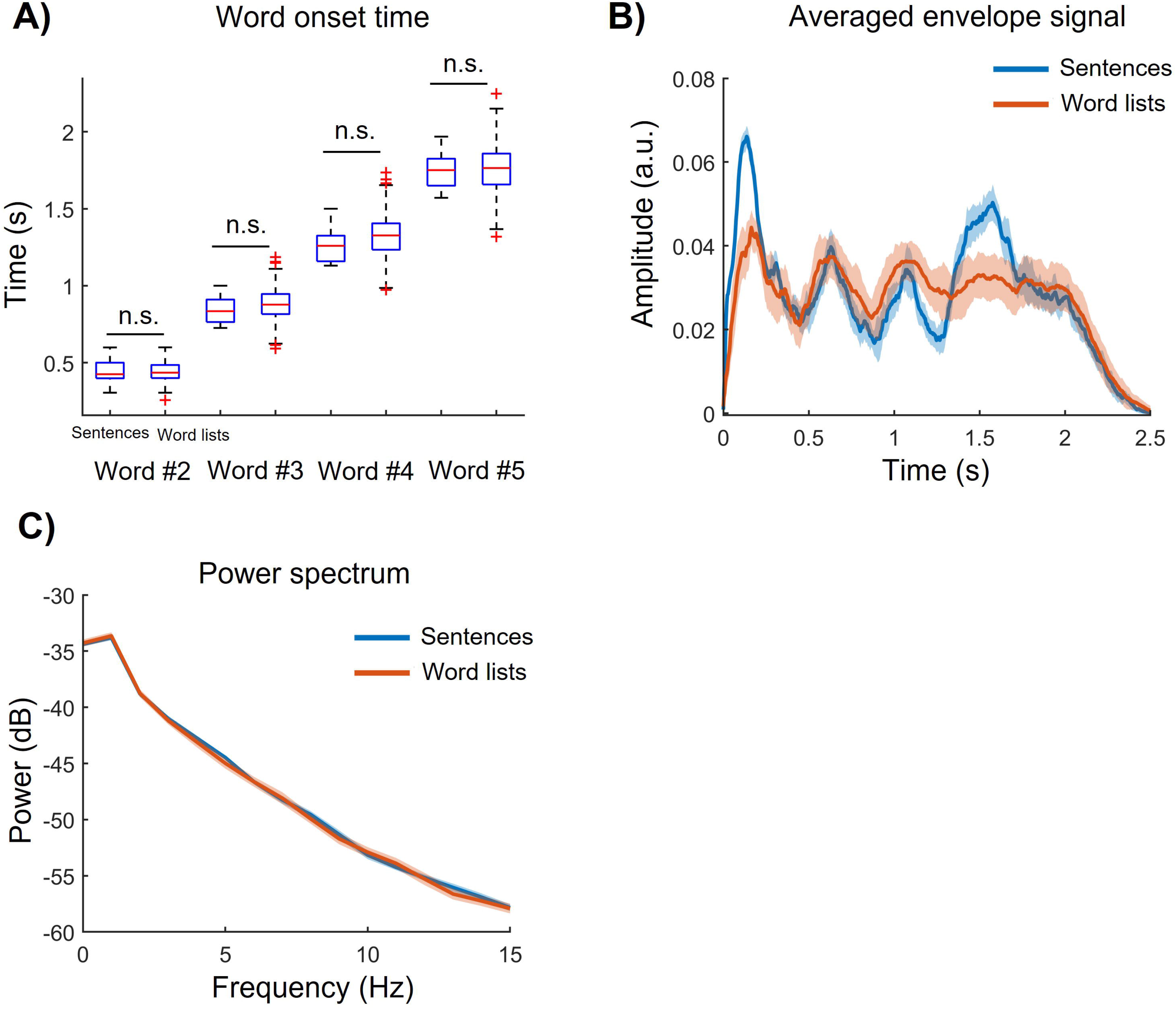
Temporal and spectral characteristics of the speech material. (A) Word onset time for each individual word in the sentences and word lists. (B) The average amplitude of the speech envelope in the sentences (blue) and word lists (red). (C) Power spectrum (0-15 Hz) of the speech envelope for the two speech types. Shaded areas around the curves reflect the standard deviation (SD).

The audio files were then masked by speech-shaped noise at SNRs of -9 dB, -6 dB, -3 dB, and 0 dB by varying the intensity of the speech while keeping the background noise constant. Speech-shaped noise was created based on the long-term power spectrum of speech. The SNR was computed from the ratio of the power of the speech signal to the power of the noise level. The intensity of the speech at the different SNRs was computed in MATLAB, and the sound was presented through MATLAB as well. The volumes were determined based on the comfort level of a few select normal-hearing participants, and none of the participants found the volume to be uncomfortable.

### 2.4. Experimental design and stimulus presentation

The experiment consisted of eight blocks, each with a randomly assigned level of SNR (-9 dB, -6 dB, -3 dB, 0 dB) and one of the two speech types (sentences or word lists). For each block, 25 trials were recorded. Each trial began with background noise, lasting 3 s plus a random interval of 0–1 s, during which subjects were asked to focus on a fixation cross on a screen in front of them. This was followed by a stimulus, in which speech was presented in the presence of background noise. After the speech presentation, the fixation cross was maintained while background noise continued for about 3 s. Finally, this was followed by a response interval in which all corpus items from the base list appeared on the screen on a 10 × 5 grid (word × category). The participants were asked to use a mouse to select the words in order that matched those they had heard. After each block (25 trials), participants were asked to rate their level of listening effort on a 1 - 10 scale using the NASA Task Load Index (Hart & Staveland, 1988) and then got a 3-minute rest.

The experiment was run using custom code in MATLAB (R2021b, MathWorks Inc.). All sounds were played through a soundcard (Scarlett 2i2 2nd Gen), and the presentation was controlled using the Psychophysics Toolbox (PTB-3). The audio signal was presented diotically through insert-earphones (a-JAYS Three). Before the main experiment, participants heard some example speech in each condition and were familiar with all procedures.

### 2.5. EEG recording and processing

The EEG data were acquired using a g.HIamp biosignal amplifier (g.tec medical engineering GmbH, Austria) with 64 channels. Electrodes were placed on a cap according to the 10-20 international system. The EEG was recorded at a sampling rate of 1,200 Hz using the left earlobe (A1) as a reference. During recordings, all electrode impedances were kept below 5 kOhm. The experiment was carried out in an electromagnetically shielded room.

The EEG data processing was carried out using a customized MATLAB script and EEGLAB toolbox (Delorme & Makeig, 2004). The data were band-passed between 0.5 Hz and 40 Hz using a 3rd-order zero-phase Butterworth filter and were then resampled to 256 Hz. Portions of data contaminated by artifacts (high-amplitude short-time activities produced by head and eye movement) were detected and corrected automatically using the Artifact Subspace Reconstruction (ASR) algorithm (Mullen et al., 2013) in EEGLAB. Independent Component Analysis (ICA) was then carried out to remove cardiac and muscle artifacts. The independent components derived by ICA were labeled using ICLabel (Pion-Tonachini et al., 2019) as implemented in EEGLAB. Components that belonged to the artifact classes (cardiac and muscle) with a probability above 50% were visually examined and intended to be removed. On average, six out of 62 components were removed per block for each participant. Three EEG channels from one participant were removed before ICA due to a high muscle artifact level and were interpolated using spline interpolation after ICA in the EEGLAB toolbox. Data were re-referenced to the average reference (Gabard-Durnam et al., 2018). For further analysis, each trial was epoched from 0 s (onset of speech) to 2 s. EEG data of two participants were excluded from further analysis due to an internal failure of the amplifier during recording.

### 2.6. Speech-brain phase locking value (PLV)

Speech stimuli were downsampled from 44.1 kHz to 1,200 Hz. The envelope of each stimulus was calculated separately in MATLAB using the Hilbert transformation. The resulting signals were downsampled to 256 Hz. Figures 2B and 2C show the average envelope signal and envelope power spectrum over trials and participants for sentences and word lists for the condition of SNR 0 dB (the mean envelope and power spectrum for all SNRs were approximately the same for each speech type).

We used the phase-locking value (PLV) (Lachaux et al., 1999), implemented in BESA Research, to quantify the phase locking between the speech envelope and the neural oscillations in different frequencies and at different time points. This measure has been used in previous studies to study cortical speech tracking (Gross et al., 2013; Molinaro et al., 2021). The PLV measures the degree to which the phase relationship between speech envelope and neural oscillations is consistent over experimental trials: 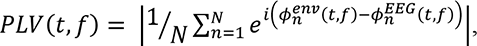 where *N* is the number of trials (here 25), *ϕ^env^*(*t*, *f*) is the phase of the speech envelope and *ϕ^EEG^*(*t*, *f*) is the phase of the EEG signal at time *t* (measured relative to the speech onset) and frequency *f*. To calculate the phase of the speech envelope and all EEG signals at each frequency, the epoched data were transformed into a time-frequency representation using a complex demodulation method implemented in BESA Research. Complex demodulation consisted of two steps. First, the time-domain signal was multiplied with a complex exponential at the frequency of interest *f*, and second, a low-pass finite impulse response (FIR) filter isolated the energy near frequency *f* (Hoechstetter et al., 2004). The data were processed for time points from 0 to 2 s post-stimulus and frequencies between 1 Hz to 20 Hz with a time-frequency sampling of 100 ms / 0.5 Hz. Furthermore, following (Peelle et al., 2013) in calculating cerebro-acoustic coherence, for each participant we used 100 random pairings of speech envelopes with EEG signal, which we average to produce random PLVs as a baseline for each condition. Then, the baseline PLVs were subtracted from the true PLVs (i.e., correct speech-brain pairing).

### 2.7. Statistical analysis

To test the effect of SNR and speech type on speech comprehension and listening effort, a two-way repeated measure analysis of variance (ANOVA) was used. Because the homogeneity of variance was violated, Greenhouse-Geisser correction was applied to ANOVA, and Wilcoxon tests were used for pairwise comparisons. All statistical analyses for behavioral data were conducted by IBM SPSS Statistics 27. For multiple testing problems, false discovery rate (FDR) correction was applied.

In order to compare the mean onset time of individual words in speech between sentences and word lists, an independent samples t-test was used. Cluster-based permutation t-test (paired, two-tailed, with 5,000 permutations, cluster entry criterion (alpha); p = 0.05) was used to test differences in the average envelope amplitude and spectrum between sentences and word lists. To compare the PLVs (averaged over different frequencies and over different points of interest) between sentences and word lists, cluster-based permutation t-tests (paired, two-tailed, with 5,000 permutations, alpha = 0.05, neighbor channel distance; 4 cm) were run for each SNR. As multiple tests were conducted, FDR-corrected p-values for each SNR were reported. To test the effect of SNR on speech-brain phase locking, PLVs of sentences and word lists were submitted to separate cluster-based permutation ANOVA tests. Cluster-based permutation tests solve the multiple comparisons problems that arise from comparing 62 electrodes and prevent inflated false-positive rates (Maris & Oostenveld, 2007). To test for a potential interaction effect of SNR and speech type, the PLVs of electrodes belonging to both significant clusters of SNR and speech type were averaged per condition and participant. On these data, we calculated a 2 (speech types) × 4 (SNR levels) repeated-measures ANOVA (as in IBM SPSS Statistics 27). BESA Statistics 2.1 was used for cluster-based permutation testing.

We investigated the association between the listening effort scores with speech-brain PLV for each speech type using a linear mixed-effect model (LMM). The fixed-effect part of the LMM consisted of SNR, speech intelligibility, and listening effort whereas the random-effect part included the variable participant.

## 3. Results

### 3.1. Behavioral performance

The mean intelligibility and mean listening effort scores are shown in Figure 3. Two-way repeated measure ANOVA was conducted to analyze potential differences across SNRs and speech types in the scores for speech intelligibility and listening effort. The results for intelligibility showed a significant interaction between speech type and SNR (*F*(3,70.4) = 13.59, *p* < 0.001, 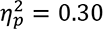), a significant main effect of speech type (*F*(1,31) = 290.60, *p* < 0.001, 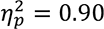), and a significant main effect of SNR (*F*(3,93) = 456.62, *p* < 0.001, 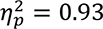).

**Figure 3.**
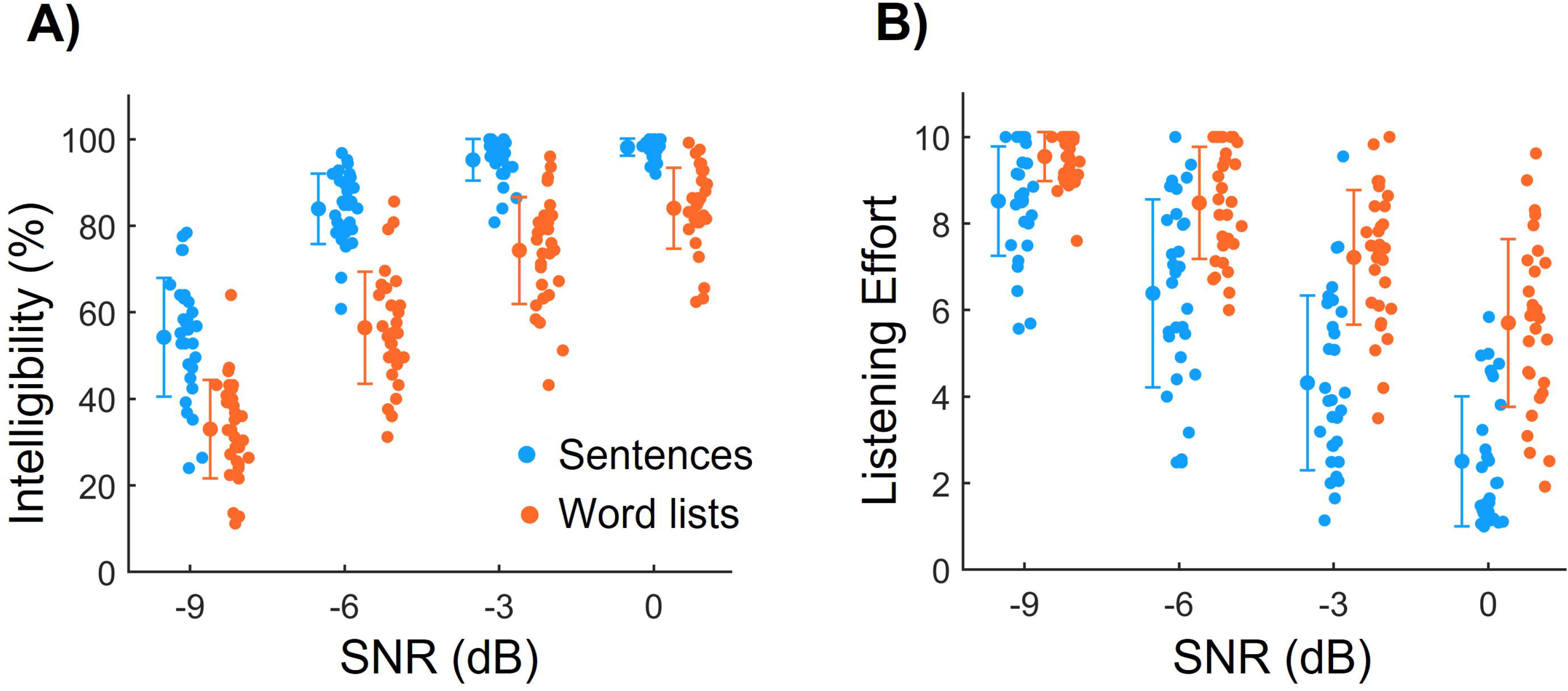
Behavioral responses. (**A**) The intelligibility of words increases with increasing SNR, both for sentences and word lists. Intelligibility is higher for sentences than for lists of random words. Dots show the results from individual subjects, and the error bars show the standard deviation. (**B**) Listening effort is higher for word lists than for sentences and decreases for both types of speech stimuli with higher SNR.

Pairwise comparisons were conducted to compare the scores for sentences with those for word lists at the same SNR level and the scores for different SNRs within each speech type. All comparisons yielded statistically significant differences (*p* < 0.001, FDR corrected), showing lower intelligibility for word lists than for sentences at the same SNR levels and increasing intelligibility for increasing SNR for both speech types. The observed interaction effect indicates the amount of decrease/increase in the intelligibility scores depends on speech types. In other words, increasing SNR from -9 dB to 0 dB was shown to increase intelligibility in sentences to the saturation level (∼ 98%). However, in word lists, this value increased only to 84%, indicating that the effect of SNR on intelligibility depends on speech type.

Similar analyses were conducted for self-reported listening efforts. The results showed a significant main interaction effect (*F*(2.47,76.8) = 12, *p* < 0.001, 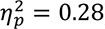), a significant effect of speech type (*F*(1,31) = 176.60, *p* < 0.001, 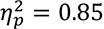), and a significant main effect of SNR (*F*(2.76,85.8) = 180, *p* < 0.001, 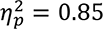). All pairwise comparisons were statistically significant (p < 0.001, FDR corrected), showing higher listening effort for word lists than for sentences at the same SNR levels and decreasing listening effort scores by increasing SNR for both speech types. The interaction effect demonstrates that the effect of SNR on listening effort scores differs across speech types. Particularly, increasing SNR by 3 dB steps from -9 dB to 0 dB decreased listening effort more for sentences than word lists at each step.

### 3.2. Phase locking between speech and brain responses

We computed PLVs between the speech envelope and the brain activity for sentences and word lists at various SNRs across different frequencies (1-20 Hz) and time points (0-2 s) (Figure 4). PLVs were averaged across the delta (1-4 Hz) and theta (4-8 Hz) frequency bands for each condition. The PLVs in the theta band showed no significant difference between conditions. Therefore, only the PLVs in the delta band were further assessed. Figure 5A shows two distinct peaks for the PLVs in the delta band, the first one from 0 ms to 500 ms and a second from 600 ms to 1100 ms.

**Figure 4.**
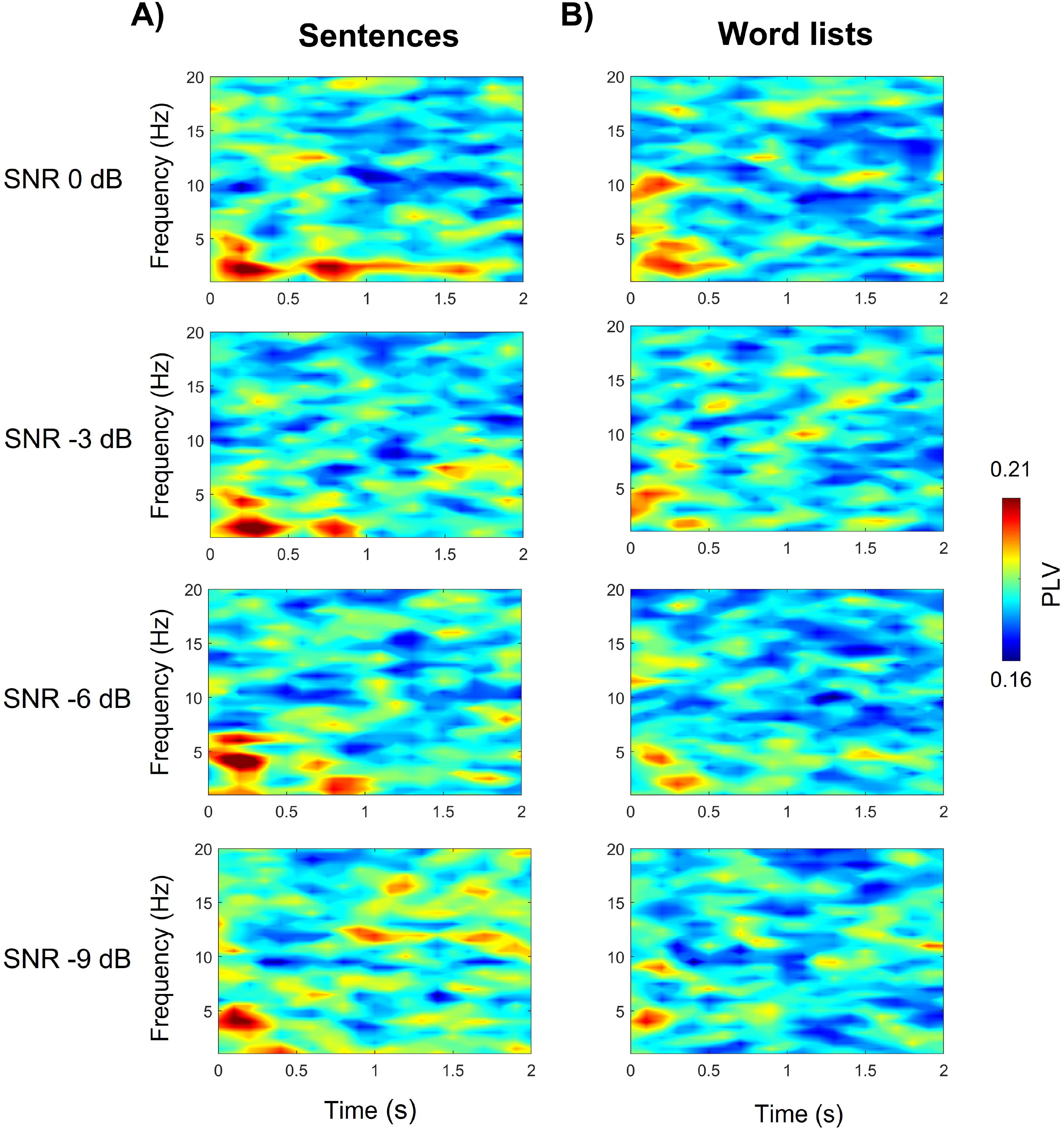
Time-frequency analysis of phase locking between the speech envelope and neural activity. The phase locking was quantified through phase-locking values (PLVs) computed for a time interval of 0-2 s and a frequency from 1 to 20 Hz. (**A**) PLVs obtained for sentences at the four signal-to-noise ratios (SNRs). (**B**) PLVss obtained for random word lists at the same SNRs.

**Figure 5.**
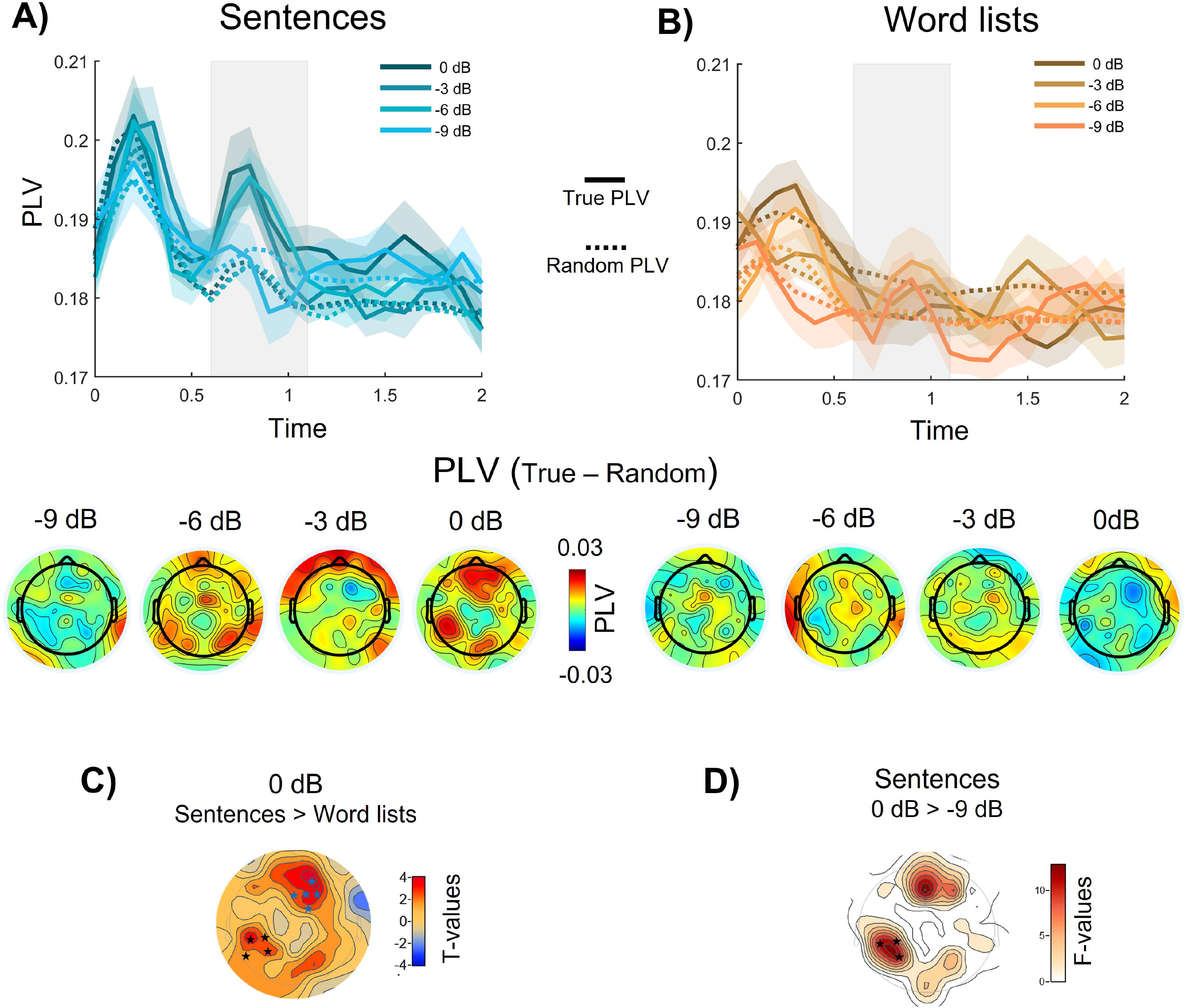
PLVs in the delta frequency band (1-4 Hz). (**A**) (top) PLVs for the correct speech-brain pairing (solid line) and random PLVs (incorrect speech-brain pairing; dashed line) obtained for sentences averaged over all 62 electrodes and over all participants for the four different signal-to-noise ratios. The gray shaded area shows a time interval of interest 600-1,100 ms. (bottom) Topographies of the PLVs (difference between true and random PLV) in the delta band averaged over the time interval of interest and all participants for different SNRs. (**B**) (top) PLVs for the speech stimuli consisting of random words averaged over electrodes and participants. (bottom) The topographies for the PLVs in the delta band averaged over the time interval of interest and participants. (C) Clusters of electrodes at which the PLVs were significantly different for the sentences and word lists at an SNR of 0 dB occurred for centro-parietal electrodes (black stars) and frontal electrodes (blue stars). (D) Statistical differences for the PLVs for sentences between the SNR of 0 dB and the SNR of -9 dB emerged at the centro-parietal electrodes (black stars; F is the F-value for the post-hoc test following the significant effect of SNR regarding PLVs for sentences). Topographies plots in panels C and D were generated in BESA Statistics 2.1.

Further, the PLVs (difference between true and random) within each of these time intervals were averaged across the time points (0-500 ms and 600-1100 ms) and were submitted to separate cluster-based t-tests to test the difference between sentences and word lists at each SNR. No significant differences were found for the first peak (*p* > 0.05) at all SNRs. For the second peak, we observed a significant difference between sentences and word lists at SNR 0 dB on frontal electrodes (AF4, Fz, F2, F4, and Fc2; p = 0.028, FDR corrected) and centro-parietal electrodes (CP5, CP3, P3, and P7; *p* = 0.035, FDR corrected) (Figure 5 C). No significant differences were found for other SNRs (*p* > 0.05).

To test the potential effect of SNR on the PLVs obtained for both sentences and word lists, the averaged PLVs were submitted to separate cluster-based permutation ANOVA tests. A significant main effect (*p* = 0.024) was obtained for sentences with a cluster of parietal electrodes (CP5, CP3, P5, P3, and PO3). No significant main effect of SNR was observed for word lists (p > 0.05). Post-hoc analysis revealed an increased PLV for sentences at 0 dB than -9 dB (*p* = 0.002) (Figure 5 D).

For testing a potential interaction effect of speech type and SNR, the PLVs of electrodes CP5, CP3, and P3, which belong to both clusters of significant differences between responses to sentences and to words lists at the SNR of 0 dB and between responses at the SNR of 0 dB and of -9 dB, were averaged (Figure 6A). The averaged PLV was submitted to two-way repeated-measures ANOVA. A significant interaction effect of speech type and SNR was found (*F*(3,87) = 8.50, *p* < 0.001, 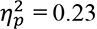). A significant main effect of speech type emerged as well (*F*(1,29) = 10.27, *p* = 0.003, 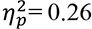), but no significant main effect of SNR (*p* > 0.05). Post-hoc analysis revealed that PLVs were higher for sentences than for word lists at SNR of 0 dB (*p* = 0.008, FDR corrected), -6 dB (*p* = 0.008, FDR corrected), and that PLVs were lower at the SNR of -9 dB than 0 dB in sentences (*p* = 0.004, FDR corrected).

**Figure 6.**
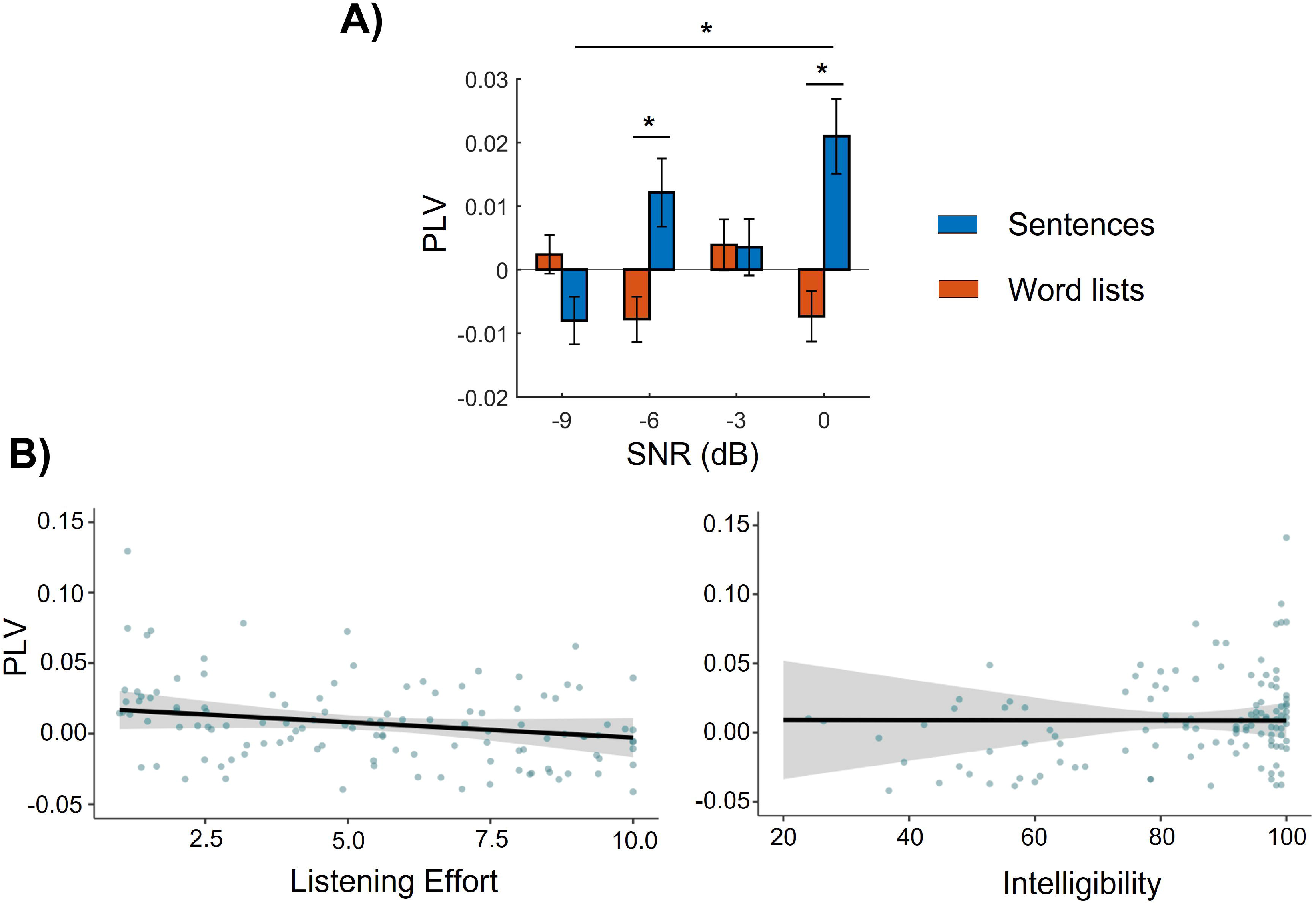
(**A**) Mean PLVs in the delta band at electrodes CP5, CP3, and P3 for responses to sentences and word lists for different SNRs. Significant differences emerge between the SNR of -9 dB and the SNR of 0 dB for responses to sentences, as well as between responses to sentences and to word lists at the SNR of -6 and 0 dB (*, p < 0.05). (**B**) Liner mixed effect model for PLVs in response to sentences. The shaded areas represent the 95% confidence interval. Results showed only a marginally significant effect of listening effort (*p* = 0.06).

To investigate the relationship between listening effort and delta PLV, we computed a LMM with SNR, listening effort, and intelligibility as the fixed-effect terms and participant as a random effect. The LMM detected a marginally significant effect of listening effort (*p* = 0.06). It further showed that SNR (*p* = 0.13) and intelligibility (*p* = 0.98) were insignificant in sentences (Figure 6B). No significant effects were found for word lists.

## 4. Discussion

This study investigated the effect of background noise and linguistic information on the phase locking between neural activity and the speech envelope in a speech-in-noise recognition task. We found significant interaction effects between the SNR and the speech type (sentences and word lists) regarding speech intelligibility, subjective listening effort, and the phase locking between delta-band neural activity and the speech envelope.

For behavioral data, we found that the effect of SNR on intelligibility and on listening effort depends on the speech type. In particular, increasing SNR increased the intelligibility score for sentences more than it did for word lists. For listening effort, increasing SNR reduced the effort more for sentences than word lists. As expected from previous studies on low- and high-predictability sentences (Bilger et al., 1984; Wilson et al., 2012), our results showed overall lower intelligibility and greater listening effort for word lists than for sentences, demonstrating the benefit of linguistic information conveyed by grammar and meaning for sentences. Indeed, the syntactic rules in sentences assist in grouping words into phrases, facilitating speech recognition and understanding (Baddeley et al., 2009; Ghitza, 2017). Semantic associations allow words to be combined into conceptual chunks and convey meaning at the sentence level (Bonhage et al., 2014). Linguistic information has accordingly been shown to reduce cognitive load in sentence recognition and memory maintenance compared to random word lists, in line with our findings (Bonhage et al., 2017).

Regarding the neural data, our analysis of the PLV between the speech envelope and the EEG recordings showed significant effects in the delta frequency band. Two distinct peaks in the PLVs at the time interval of 0-500 ms and 600-1,100 ms after speech onset were observed for sentences. The first peak that emerged for both true and random PLVs may be an evoked response to the onset of speech. Therefore, no differences between sentences and word lists were observed for this peak. Nonetheless, significantly higher speech-brain phase locking in the time interval of 600-1100 ms was observed for sentences than for word lists at the SNR of 0 and -6 dB. This might be related to the top-down control of the phase locking by high-level linguistic processing. Sentences were made up of subject, verb, numeral, adjective, and object structure. The increased speech-brain phase locking for sentences compared to word lists starts approximately 500 ms after speech onset which is comparable to the onset of the second word (verb) of the sentences which is about 451 ms (comparably, in word lists, the second word occurs approx 443 ms after speech onset but did not yield an elevated PLV).

The increased PLV at this latency for sentences might be related to the cortical tracking of subject-verb structures in a sentence reported by Ding et al. (2015). They demonstrated that the cortical response gradually decreased within the noun phrase, then showed a transient increase after the onset of the verb phrase. This indicates that cortical activity is entrained into linguistic structures that are constructed internally, based on syntax (Peelle et al., 2013; Ding et al., 2015). Indeed other studies also showed delta-band speech tracking relates to the encoding of syntactic information in connected speech (Meyer & Gumbert, 2018; Molinaro & Lizarazu, 2018). For example, Lu et al. (2022) and Coopmans et al. (2022) studied the influence of sentential structure on neural tracking of word sequences and found a significantly stronger delta-band neural response to regular sentences than to word lists, suggesting that delta-band neural responses are modulated by the compositional meaning of sentence structures.

The interaction between SNR and speech type was characterized by the different effects of SNR on the PLV in response to sentences and word lists. In sentences, greater PLV values were observed at the SNR of 0 dB than at the SNR of -9 dB. The coupling between brain activity and speech is presumably reduced by noise through energetic masking (Dimitrijevic et al., 2019). Speech-shaped noise matches the long-term spectral properties of the speech signal, and the spectrotemporal energies overlap in a combination of speech and noise. When the SNR is low, the amplitude (energy) of the noise will dominate the neural representation in the auditory nervous system, resulting in the poor neural representation of the target signal (Wang & Xu, 2021; Brungart, 2001).

Our LMM model results showed that the lowest p-value emerged for the listening effort, indicating a trend for PLV to reflect the listening effort, although this variable was not statistically significant. Despite the lack of significance, this trend corroborates with the literature (Decruy et al., 2020; Dimitrijevic et al., 2019), showing that speech tracking decreases with listening effort.

In this study, the experimental conditions were presented using a block design paradigm. This paradigm allows manipulating task demands across blocks to identify neural responses associated with specific processes, but it also has some disadvantages. In each block, participants could anticipate the type of stimulus (sentences and random word lists) and the level of SNR in subsequent trials. Therefore, they might systematically change their level of attention and effort across conditions, impacting both the behavioral and the neural data (Humphries et al. 2006).

### Data availability statement

The EEG data, behavioral data, and audio files can be download on OSF (https://osf.io/b9wdp/?view_only=0fd2608f0ca7437aa633e67d2c412744).

## 5. Conclusion

In this study, we examined the effect of background noise level and linguistic information on speech-brain coupling during speech-in-noise recognition. Results showed an interaction between SNR and linguistic information on the phase locking between neural activity in the delta-band and the amplitude modulation in speech. Increased PLVs for sentences as compared to random word lists were observed, indicating that linguistic structure increases the PLVs. A decrease in PLVs at the SNR of -9 dB compared to the SNR of 0 dB in sentences emerged as well, likely indicating a disruption in acoustic properties by energetic masking that reduces speech tracking. Last but not least, the PLV for sentences showed a decreasing trend with increasing listening effort, indicating that listening effort might be decoded from brain activity, although the trend needs to be substantiated with larger-scale studies.

## Notes

https://osf.io/b9wdp/?view_only=0fd2608f0ca7437aa633e67d2c412744

### Competing Interest Statement

The authors have declared no competing interest.

## References

Baddeley, A. D., Hitch, G. J., & Allen, R. J. (2009). Working memory and binding in sentence recall. Journal of Memory and Language, 61(3), 438–456.

Bilger, R. C., Nuetzel, J. M., Rabinowitz, W. M., & Rzeczkowski, C. (1984). Standardization of a Test of Speech Perception in Noise. Journal of Speech and Hearing Research, 27(1), 32–48.

Bonhage, C. E., Fiebach, C. J., Bahlmann, J., & Mueller, J. L. (2014). Brain signature of working memory for sentence structure: enriched encoding and facilitated maintenance. Journal of Cognitive Neuroscience, 26(8), 1654–1671.

Bonhage, C. E., Meyer, L., Gruber, T., Friederici, A. D., & Mueller, J. L. (2017). Oscillatory EEG dynamics underlying automatic chunking during sentence processing. NeuroImage, 152, 647–657.

Coopmans, C. W., de Hoop, H., Hagoort, P., & Martin, A. E. (2022). Effects of Structure and Meaning on Cortical Tracking of Linguistic Units in Naturalistic Speech. Neurobiology of Language, 3(3), 386–412.

Dai, B., McQueen, J. M., Terporten, R., Hagoort, P., & Kösem, A. (2022). Distracting linguistic information impairs neural tracking of attended speech. Current Research in Neurobiology, 3, 100043.

Decruy, L., Lesenfants, D., Vanthornhout, J., & Francart, T. (2020). Top-down modulation of neural envelope tracking: The interplay with behavioral, self-report and neural measures of listening effort. European Journal of Neuroscience, 52(5), 3375–3393.

Delorme, A., & Makeig, S. (2004). EEGLAB: an open source toolbox for analysis of single-trial EEG dynamics including independent component analysis. Journal of Neuroscience Methods, 134(1), 9–21.

Dimitrijevic, A., Smith, M. L., Kadis, D. S., & Moore, D. R. (2019). Neural indices of listening effort in noisy environments. Scientific Reports, 9(1), 11278.

Ding, N., Melloni, L., Zhang, H., Tian, X., & Poeppel, D. (2015). Cortical tracking of hierarchical linguistic structures in connected speech. Nature Neuroscience 2016 19:1, 19(1), 158–164.

Ding, N., & Simon, J. Z. (2013). Adaptive Temporal Encoding Leads to a Background-Insensitive Cortical Representation of Speech. The Journal of Neuroscience, 33(13), 5728.

Gabard-Durnam, L. J., Leal, A. S. M., Wilkinson, C. L., & Levin, A. R. (2018). The harvard automated processing pipeline for electroencephalography (HAPPE): Standardized processing software for developmental and high-artifact data. Frontiers in Neuroscience, 12, 97.

Ghinst, M. vander, Bourguignon, M., Niesen, M., Wens, V., Hassid, S., Choufani, G., Jousmäki, V., Hari, R., Goldman, S., & de Tiège, X. (2019). Cortical Tracking of Speech-in-Noise Develops from Childhood to Adulthood. Journal of Neuroscience, 39(15), 2938–2950.

Ghitza, O. (2017). Acoustic-driven delta rhythms as prosodic markers. Language, Cognition and Neuroscience, 32(5), 545– 561.

Giraud, A. L., & Poeppel, D. (2012). Cortical oscillations and speech processing: emerging computational principles and operations. Nature Neuroscience 2012 15:4, 15(4), 511–517.

Gross, J., Hoogenboom, N., Thut, G., Schyns, P., Panzeri, S., Belin, P., & Garrod, S. (2013). Speech Rhythms and Multiplexed Oscillatory Sensory Coding in the Human Brain. PLOS Biology, 11(12), e1001752.

Hart, S. G., & Staveland, L. E. (1988). Development of NASA-TLX (Task Load Index): Results of Empirical and Theoretical Research. Advances in Psychology, 52(C), 139–183.

Hoechstetter, K., Bornfleth, H., Weckesser, D., Ille, N., Berg, P., & Scherg, M. (2004). BESA source coherence: a new method to study cortical oscillatory coupling. Brain Topography, 16(4), 233–238.

Kerlin, J. R., Shahin, A. J., & Miller, L. M. (2010). Attentional Gain Control of Ongoing Cortical Speech Representations in a “Cocktail Party.” Journal of Neuroscience, 30(2), 620–628.

Keshavarzi, M., Kegler, M., Kadir, S., & Reichenbach, T. (2020). Transcranial alternating current stimulation in the theta band but not in the delta band modulates the comprehension of naturalistic speech in noise. NeuroImage, 210, 116557.

Kolozsvári, O. B., Xu, W., Gerike, G., Parviainen, T., Nieminen, L., Noiray, A., & Hämäläinen, J. A. (2021). Coherence Between Brain Activation and Speech Envelope at Word and Sentence Levels Showed Age-Related Differences in Low Frequency Bands. Neurobiology of Language, 2(2), 226–253.

Lachaux, J.-P., Rodriguez, E., Martinerie, J., & Varela, F. J. (1999). Measuring Phase Synchrony in Brain Signals. Hum Brain Mapping, 8, 194–208.

Lu, Y., Jin, P., Pan, X., & Ding, N. (2022). Delta-band neural activity primarily tracks sentences instead of semantic properties of words. NeuroImage, 251, 118979.

Maris, E., & Oostenveld, R. (2007). Nonparametric statistical testing of EEG- and MEG-data. Journal of Neuroscience Methods, 164(1), 177–190.

McHaney, J. R., Gnanateja, G. N., Smayda, K. E., Zinszer, B. D., & Chandrasekaran, B. (2021). Cortical Tracking of Speech in Delta Band Relates to Individual Differences in Speech in Noise Comprehension in Older Adults. Ear and Hearing, 343–354.

Meyer, L., & Gumbert, M. (2018). Synchronization of Electrophysiological Responses with Speech Benefits Syntactic Information Processing. Journal of Cognitive Neuroscience, 30(8), 1066–1074.

Molinaro, N., & Lizarazu, M. (2018). Delta(but not theta)-band cortical entrainment involves speech-specific processing. European Journal of Neuroscience, 48(7), 2642–2650.

Molinaro, N., Lizarazu, M., Baldin, V., Pérez-Navarro, J., Lallier, M., & Ríos-López, P. (2021). Speech-brain phase coupling is enhanced in low contextual semantic predictability conditions. Neuropsychologia, 156.

Mohammadi, Y., Graversen, C., Manresa, J. B., Østergaard, J., Andersen, O. K. (2023). Effects of background noise and linguistic violations on frontal theta oscillations during effortful listening. “In review”.

Mullen, T., Kothe, C., Chi, Y. M., Ojeda, A., Kerth, T., Makeig, S., Cauwenberghs, G., & Jung, T. P. (2013). Real-time modeling and 3D visualization of source dynamics and connectivity using wearable EEG. Annual International Conference of the IEEE Engineering in Medicine and Biology Society. IEEE Engineering in Medicine and Biology Society. Annual International Conference, 2013, 2184–2187.

O’Sullivan, J. A., Power, A. J., Mesgarani, N., Rajaram, S., Foxe, J. J., Shinn-Cunningham, B. G., Slaney, M., Shamma, S. A., & Lalor, E. C. (2015). Attentional Selection in a Cocktail Party Environment Can Be Decoded from Single-Trial EEG. Cerebral Cortex, 25(7), 1697–1706.

Pasley, B. N., David, S. v., Mesgarani, N., Flinker, A., Shamma, S. A., Crone, N. E., Knight, R. T., & Chang, E. F. (2012). Reconstructing Speech from Human Auditory Cortex. PLOS Biology, 10(1), e1001251.

Peelle, J. E., Gross, J., & Davis, M. H. (2013). Phase-locked responses to speech in human auditory cortex are enhanced during comprehension. Cerebral Cortex (New York, N.Y.: 1991), 23(6), 1378–1387.

Petersen, E. B., Wöstmann, M., Obleser, J., & Lunner, T. (2017). Neural tracking of attended versus ignored speech is differentially affected by hearing loss. Journal of Neurophysiology, 117(1), 18–27.

Pion-Tonachini, L., Kreutz-Delgado, K., & Makeig, S. (2019). ICLabel: An automated electroencephalographic independent component classifier, dataset, and website. NeuroImage, 198, 181–197.

Riecke, L., Formisano, E., Sorger, B., Başkent, D., & Gaudrain, E. (2018). Neural Entrainment to Speech Modulates Speech Intelligibility. Current Biology, 28(2), 161–169.e5.

Rimmele, J. M., Zion Golumbic, E., Schröger, E., & Poeppel, D. (2015). The effects of selective attention and speech acoustics on neural speech-tracking in a multi-talker scene. Cortex; a Journal Devoted to the Study of the Nervous System and Behavior, 68, 144–154.

vander Ghinst, M., Bourguignon, M., op de Beeck, M., Wens, V., Marty, B., Hassid, S., Choufani, G., Jousmäki, V., Hari, R., van Bogaert, P., Goldman, S., & de Tiège, X. (2016). Left Superior Temporal Gyrus Is Coupled to Attended Speech in a Cocktail-Party Auditory Scene. Journal of Neuroscience, 36(5), 1596–1606.

Wang, X., & Xu, L. (2021). Speech perception in noise: Masking and unmasking. Journal of Otology, 16(2), 109–119.

Wilsch, A., Neuling, T., Obleser, J., & Herrmann, C. S. (2018). Transcranial alternating current stimulation with speech envelopes modulates speech comprehension. NeuroImage, 172, 766–774.

Wilson, R. H., McArdl, R., Watt, K. L., & Smith, S. L. (2012). The revised speech perception in noise test (R-SPIN) in a multiple signal-to-noise ratio paradigm. Journal of the American Academy of Audiology, 23(8), 590–605.

Zoefel, B., Archer-Boyd, A., & Davis, M. H. (2018). Phase Entrainment of Brain Oscillations Causally Modulates Neural Responses to Intelligible Speech. Current Biology, 28(3), 401–408.e5.

